# Brain-like learning with exponentiated gradients

**DOI:** 10.1101/2024.10.25.620272

**Authors:** Jonathan Cornford, Roman Pogodin, Arna Ghosh, Kaiwen Sheng, Brendan A. Bicknell, Olivier Codol, Beverley A. Clark, Guillaume Lajoie, Blake A. Richards

## Abstract

Computational neuroscience relies on gradient descent (GD) for training artificial neural network (ANN) models of the brain. The advantage of GD is that it is effective at learning difficult tasks. However, it produces ANNs that are a poor phenomenological fit to biology, making them less relevant as models of the brain. Specifically, it violates Dale’s law, by allowing synapses to change from excitatory to inhibitory, and leads to synaptic weights that are not log-normally distributed, contradicting experimental data. Here, starting from first principles of optimisation theory, we present an alternative learning algorithm, exponentiated gradient (EG), that respects Dale’s Law and produces log-normal weights, without losing the power of learning with gradients. We also show that in biologically relevant settings EG outperforms GD, including learning from sparsely relevant signals and dealing with synaptic pruning. Altogether, our results show that EG is a superior learning algorithm for modelling the brain with ANNs.

## Introduction

Computational neuroscience has recently become more concerned with building models that can learn ethological behaviours from realistic, high-dimensional inputs (Summerfield, 2022). This is important for the role of computational models as the interface between theory and empirical data (Levenstein et al., 2023), since models that can learn more realistic tasks can be directly compared to experimental data (Doerig et al., 2023). For this purpose, gradient descent (GD) is increasingly used in computational neuroscience studies (Costa et al., 2017; Driscoll et al., 2024; Flesch et al., 2022; Langdon and Engel, 2022; Malkin et al., 2024; Tang et al., 2023; Whittington et al., 2020; Yamins and DiCarlo, 2016; Yang et al., 2019). GD is leveraged in these studies because it is very effective for training artificial neural networks (ANNs), even though it is not considered biologically plausible (Lillicrap et al., 2020). Moreover, it is clear that GD is phenomenologically different from the learning algorithm(s) of the brain. First, GD prescribes synaptic changes that violate Dale’s law, i.e. changes from synaptic inhibition to excitation (Eccles, 1976). Second, GD does not produce synaptic weights that are log-normal (Pogodin et al., 2024), which have been observed across species and brain regions (Buzsáki and Mizuseki, 2014; Dorkenwald et al., 2022; Rößler et al., 2023; Song et al., 2005). Even if we set aside concerns about the biological plausibility of GD, these phenomenological differences motivate the search for an alternative more brain-like learning algorithm for training models of neural circuits. However, for an alternative algorithm to be better than GD for computational neuroscience, it should be no less powerful at learning than GD, and ideally better at learning in biologically relevant situations.

GD is effective at learning difficult tasks because it is fundamentally normative (Bredenberg and Savin, 2023; Levenstein et al., 2023). Specifically, GD provides the greatest improvement in a measure of behavioural performance with the smallest change to synapses as possible (if we restrict ourselves to linear estimates of performance). But, what does it mean to measure synaptic weight change? Here we re-analyse this core theoretical assumption and show that GD does not align with biology, in part, because it is based on a measure of synaptic change that is agnostic to changes between inhibition and excitation. Using this insight we explore a related, but different learning algorithm, exponentiated gradient (EG) (Kivinen and Warmuth, 1997). Like GD, EG is also derived from normative first principles, and leverages gradient signals for learning, but it is based on a measure of synaptic change that adheres to Dale’s law. Crucially, it works as well as GD for deep learning (Bernstein et al., 2020; Schwarz et al., 2021; Sun et al., 2022), which rules out the common concern that brain-like algorithms do not scale to hard tasks (Bartunov et al., 2018).

We show that EG has a number of desirable properties for modelling the brain. First, by construction, EG maintains Dale’s law. Second, EG leads to networks with heavy-tailed, log-normal distributions of synaptic weights. Third, synaptic changes become proportional to existing synaptic weights, as has been observed experimentally (Loewenstein et al., 2011; Melander et al., 2021). Fourth, in line with EG being a better algorithm for modelling real neural learning, we find that neural networks trained with EG perform better than their GD trained counterparts in biologically relevant learning situations. For example, we find EG works better with synaptic pruning (Faust et al., 2021; Maret et al., 2011) and in situations where there are irrelevant inputs, as is common in the brain. Specifically, with EG, multi-compartmental neuron models better learn to identify task-relevant synapses, and likewise recurrent neural networks (RNNs) are better at sensorimotor tasks in the presence of many irrelevant signals. Altogether, these results show that EG is as or more powerful for learning as GD in biologically relevant contexts. As well, it provides a better phenomenological match to biological learning. This indicates that EG is a better learning algorithm than GD for modelling the brain.

## Results

### Deriving learning algorithms from optimisation theory

We begin with a brief discussion of how GD and EG can be derived from first principles, as this can explain why they are both effective learning algorithms, and illustrate why EG naturally respects Dale’s law.

Both GD and EG can be derived by framing the problem of learning as follows: how can we obtain the greatest improvement in performance with the least amount of synaptic change? This question can be addressed with the mirror descent framework from machine learning (Bubeck, 2015; Nemirovskij and Yudin, 1983). According to the mirror descent framework, changes to the synaptic weights in a neural network should minimise a combination of task error (quantified by a loss function) and a “synaptic change penalty”. This is formalised as:

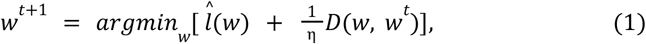

where *w*^*t*^ are the synaptic weights at time 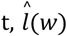 is a linear approximation of the loss function *l*(*w*) around *w*^*t*^, *D*(*w, w*^*t*^) is the synaptic change penalty from *w* to a candidate *w*, and η defines the strength of the synaptic change penalty, thereby controlling the trade-off between reducing the loss and keeping weight changes small. Note that η will later play the role of the learning rate (Eqs. 2, 5).

The form that *D* takes will lead to different learning algorithms. Notably, GD follows from measuring synaptic change with the squared 2-norm, i.e. the squared Euclidean distance between the two synaptic weight vectors (Fig. 1A, left):

**Figure 1:**
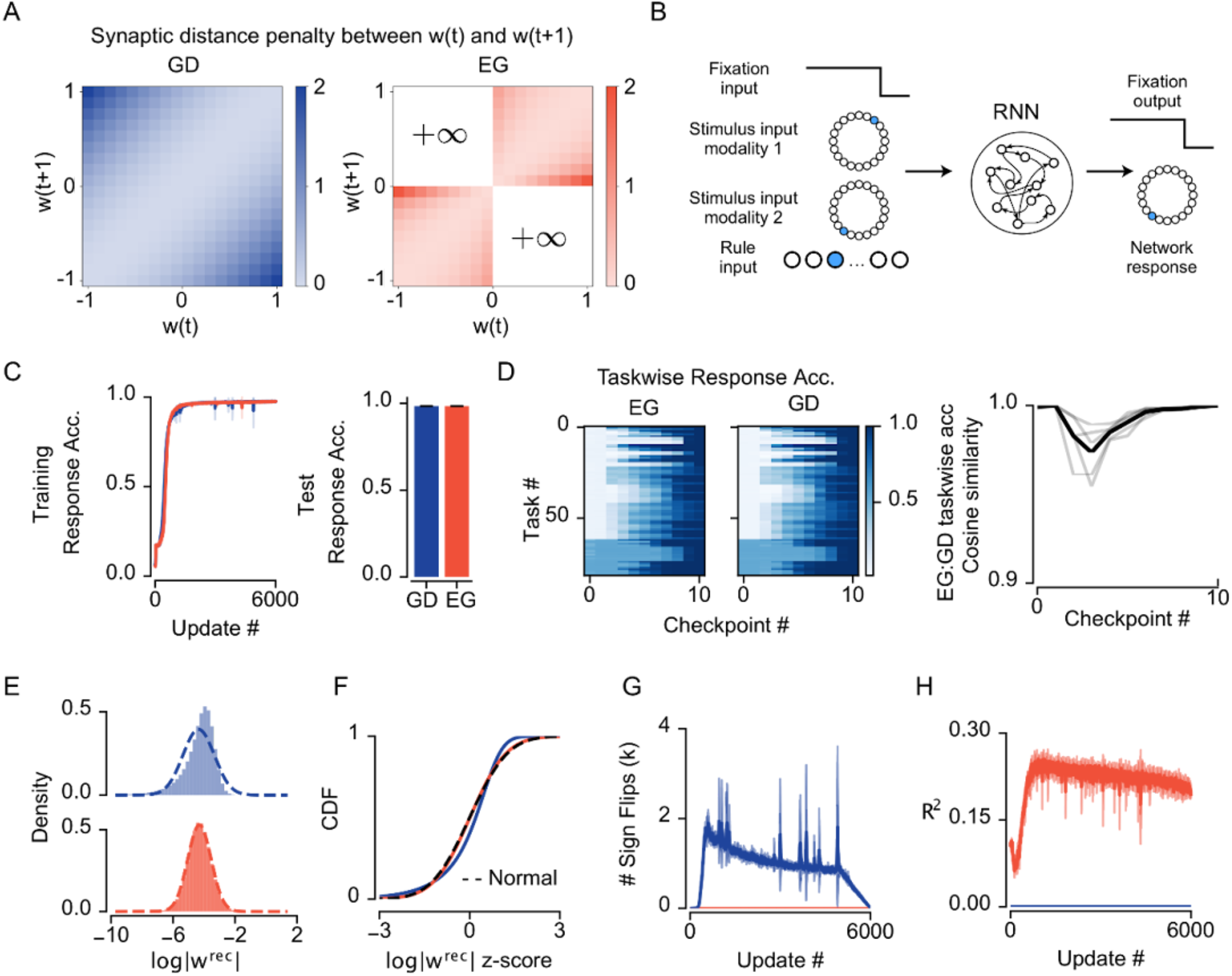
Exponentiated gradient (EG) solves the Mod-Cog suite of tasks as well as gradient descent (GD), but with more biological weight solutions. **a**| Left: GD weight geometry is Euclidean and treats E/I weights equally. Right: EG weight geometry is non-Euclidean and prohibits switching between E and I weights. **b**| Schematic of the Mod-Cog suite of tasks solved by a recurrent neural network (RNN); adapted from (Khona et al., 2023). **c**| Accuracy of EG and GD trained networks during learning (n=5). Response denotes the time period when the fixation input drops to 0. **d**| Left: taskwise testset performance of networks at different checkpoints during training. Checkpoints were taken at updates #50, #100, and at every 10% performance threshold from 25%-95%, and at the end of training. Right: cosine similarity (see methods) of taskwise accuracy between EG and GD trained network checkpoints (black: mean, grey: pairs of experiments). **e**| Histogram of final recurrent weights for GD (top) and EG (bottom). Dashed line shows a normal pdf for comparison. **f**| Same as **e**, but for CDFs rather than densities. **g**| Number of synaptic weight sign flips per update during training (dark: mean, light: std. deviation, n=5). **h**| R^2 of a linear fit to update size from current weight (n=5).

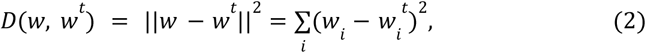

where *i* is an index over individual synaptic weights, and *w* is the candidate for new synaptic weights. If we solve equation (1), given equation (2), we obtain GD:

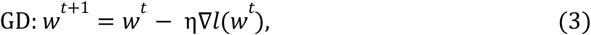

where ∇*l* indicates the gradient of the loss function with respect to the synaptic weights, and η indicates the learning rate (see appendix A for details of this derivation).

Note, however, that equation (2) makes no distinction between changes to synapses that obey Dale’s law and those that don’t (i.e. it doesn’t distinguish between changes that preserve the sign of synapses and those that don’t). An alternative choice for *D* that does respect this fundamental constraint is the unnormalised relative entropy (Fig. 1A, right), given by:

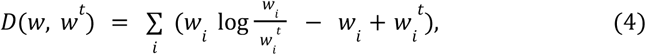

where we have assumed positive weights for now. If we consider a scenario where a candidate weight, *w*_*j*_, has a different sign than the current weight, 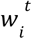, then equation (4) is effectively an infinite penalty since the log is undefined. Put another way, this version of *D* simply does not allow for synapses to change from excitatory to inhibitory. Therefore, when solving equation (1), given equation (4), the resulting weights do not change signs. If we incorporate negative weights into the architecture, we get EG (Kivinen and Warmuth, 1997):

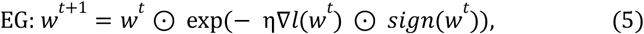

where ⊙ denotes element-wise multiplication. Unlike GD, EG does not treat all synaptic weight changes as equal – it instead favours changing already large synapses and completely prevents synapses from changing from inhibitory to excitatory, or vice-versa (because the exponential is always positive, the update can only scale the magnitude of the weight). Not only do these properties respect Dale’s law, they also lead to log-normal synaptic weights, as we show below.

### EG is a better phenomenological fit to biology than GD

To investigate the impact of using either GD or EG for learning, we performed experiments with RNNs learning a suite of cognitive neuroscience style tasks named Mod-Cog (Khona et al., 2023; Molano-Mazon et al., 2022; Yang et al., 2019). We chose these tasks because they are a representative sample of the tasks often used in behavioural neuroscience experiments and they have provided fruitful insights into the nature of learning in RNNs in previous studies (Driscoll et al., 2024; Yang et al., 2019).

In the Mod-Cog tasks, an RNN receives four inputs: a fixation cue, a task cue, and two input variables each encoding a circular position. Depending on the task given (as indicated by the task cue) and the time varying input representations, the network must give a specific output. For example, in one task, the network must simply remember the value of one of the input variables and output that value during the response period, denoted by when the fixation cue drops to 0 (Fig. 1B, S1). In other more complicated tasks, the network must estimate and calculate output sequences using delays between inputs. Overall, there were 82 such tasks, each with a different logic for producing the correct output. Networks were trained on all tasks simultaneously and tested on a subset of held out data (see Methods for more details).

Networks trained with either GD or EG learned to perform all tasks with high accuracy. Learning with both algorithms converged relatively rapidly, reaching almost peak performance after 1000 weight updates (Fig. 1C, left), with a final accuracy of 98.4% for both GD and EG (Fig. 1C, right). This indicates that both algorithms are effective for training RNNs on these simple cognitive tasks.

We next asked whether the algorithms found different solutions, despite being equally good at learning. To start, we analysed the distribution and spectral properties of the network activity when solving the tasks, but found little difference between networks trained by EG or GD (Fig. S2). Turning next to learning trajectories, we explored whether the two algorithms learned the tasks in a different order, which would indicate different solutions because tasks were easier or harder for the two algorithms. To examine this we took checkpoints of the models throughout training and measured the test accuracy on each of the tasks (Fig. 1D, left). When we compared the two algorithms’ profiles across the tasks at each of the checkpoint (measured as the cosine similarity between the two vectors of per task accuracies), we found that the two algorithms largely learned the tasks in the same order, indicating that the difficulty of each task was roughly equivalent for the two algorithms (Fig. 1D, right).

The final synaptic weight distributions discovered by the two algorithms, however, were different. We initialised the networks from random log-normal weight distributions (see Methods), because experimental data shows that brains typically have log-normal distributions in their synaptic strengths (as measured by electrophysiology or spine size) (Buzsáki and Mizuseki, 2014; Dorkenwald et al., 2022; Rößler et al., 2023; Song et al., 2005). When we examined the distribution of synaptic weights after training, we found that the weights of networks trained with GD were no longer log-normal (Fig. 1E, top). In contrast, weights in networks trained with EG were still well fit by a log-normal distribution after training (Fig. 1E, bottom). As a result, the value of the Kolmogorov-Smirnov (see Methods) statistic for the GD distribution was 10 times larger than that for the EG distribution, and the two distributions were different from each other (Fig. 1F, Fig. S3B). Furthermore, starting from initial distributions that were normal, uniform or log-normal, we consistently found that the changes prescribed by EG were approximately log-normal, whereas instead GD changes were approximately normal (Fig. S3). Thus, GD and EG discover different synaptic weight solutions, despite exhibiting similar levels of accuracy across the tasks.

We then explored the implications with respect to Dale’s law, which implies that synapses cannot change from being excitatory to being inhibitory, and vice-versa (Eccles, 1976). Notably, GD does not provide any prohibition against changing the sign of a synaptic weight between positive or negative, whereas EG prevents such sign changes explicitly. Thus, if a network is initialised to obey Dale’s law, GD could alter the identities of excitatory and inhibitory neurons, whereas EG cannot. To confirm this, we examined the number of synaptic sign flips that occurred in the networks trained with GD and EG, and as expected, GD exhibited many synaptic sign flips throughout learning whereas EG exhibited none (Fig. 1G). Hence, EG relies on weight space updates that respect Dale’s law to find a solution, whereas GD does not.

Finally, we were interested in exploring how well the learning dynamics of the two algorithms matched previously observed neuroscientific studies. One notable, and replicated, observation has been the magnitude of weight updates, i.e. the amount of synaptic change that occurs, is proportional to the current synaptic weight magnitude (Loewenstein et al., 2011; Melander et al., 2021). Considering the multiplicative form of the EG update rule (Eq. 5), we expect EG to produce proportional weight updates and GD to not. This analysis, however, assumes that learning signals are independent from the synaptic weight magnitudes, which is not necessarily true in RNNs. To test this empirically, we fit a linear regression model from the absolute value of the synaptic weights to the magnitude of synaptic plasticity for each weight update. Here we found that for GD this model did not explain any of the update variance, whereas for EG a reasonable proportion of the variance was explained (Fig. 2H). As such, we conclude that EG synaptic changes are proportional, which matches known biology, whereas GD synaptic changes are not.

**Figure 2:**
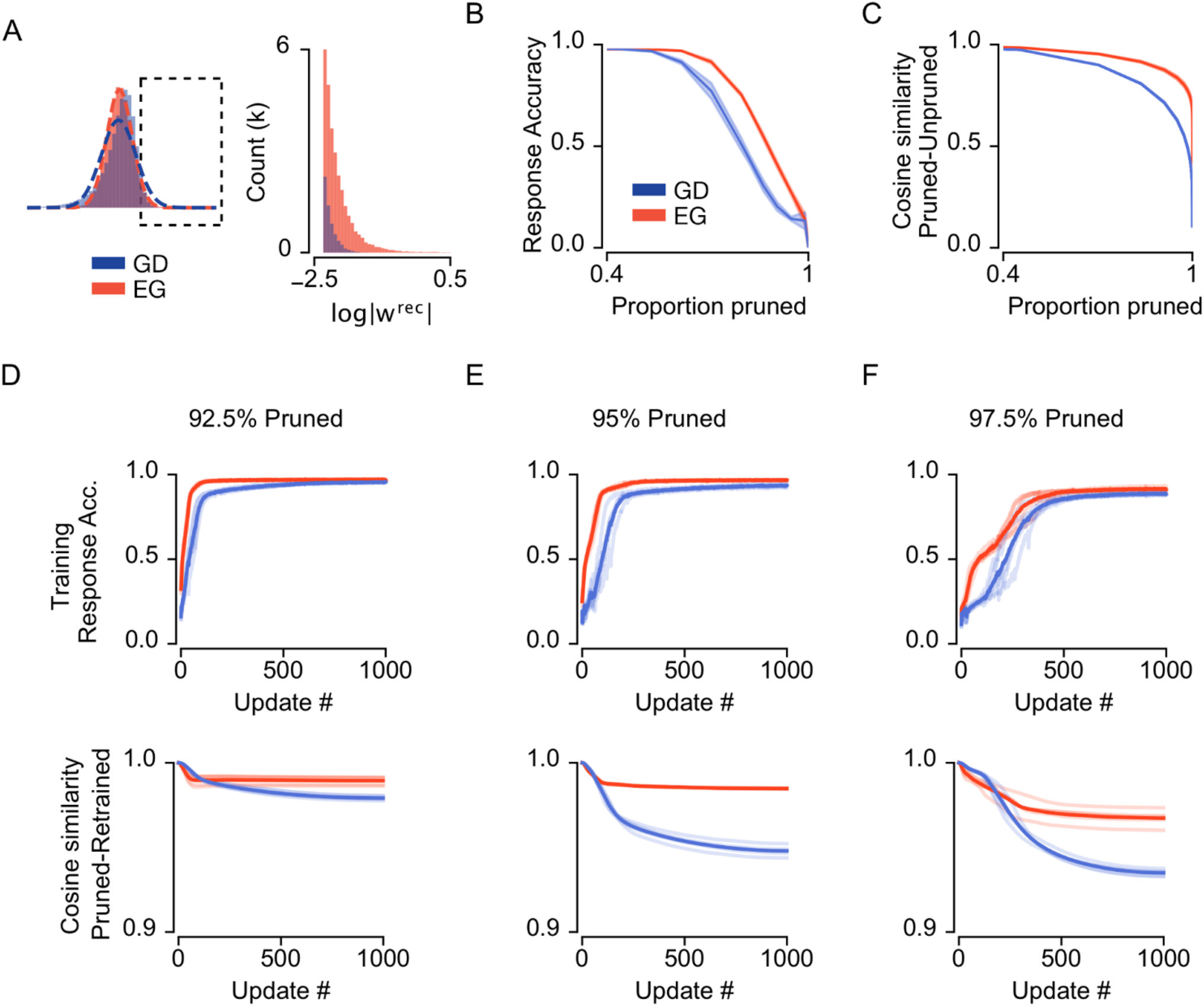
Exponentiated gradient (EG) finds weights that are easier to prune, and easier to re-train after pruning, than gradient descent (GD). **a**| The right tail of the final weight distributions for EG (orange) contains many more weights than for GD (same data as Fig. 1, k : thousand) **b**| Response accuracy for networks (from Fig. 1) as smaller weights are pruned from the network. **c**| Cosine similarity between pruned and unpruned networks as smaller weights are pruned from the network. **d**| Top: accuracy of 92.5% pruned network re-learning Mod-Cog tasks. Bottom: cosine similarity between the retrained network and the original pruned network (from **b**). **e**| As **d**, but for 95% pruning level. **f**| As **d**, but for 97.5% pruning level. (n=5, EG orange, GD blue)

In sum, these results demonstrate that both GD and EG are effective learning algorithms for training RNNs on cognitive tasks, but EG produces solutions that are a better fit to existing neuroscience data: they match the distribution of synaptic weights seen in real brains, they respect the constraints of Dale’s law, and they exhibit proportionality of weight updates. Altogether, this demonstrates that training with EG provides a better fit to biology than using training with GD.

### Learning with EG is more robust to synaptic pruning

Having observed that the EG algorithm provides a better phenomenological fit to existing biological data than GD, we next tested whether EG is also a better learning algorithm for biologically-relevant situations. A notable difference between the two algorithms is the skewed, heavier tailed synaptic weight distributions found by EG (Fig. 1E, 2A). This observation motivated us to investigate how robust the EG and GD trained networks are to synaptic pruning, an important cellular process that occurs during sleep and development (Faust et al., 2021; Maret et al., 2011). We hypothesised the large weights found by EG form a backbone that dominates computational dynamics (Teramae and Fukai, 2014), and thus, pruning the smaller weights would have relatively little effect on performance compared to GD trained networks.

To test network robustness to pruning, we took the networks trained on Mod-Cog and examined their accuracy after removing an increasing proportion of the smallest weights. We found that the EG trained networks maintained accuracy in the face of synaptic pruning much better than the GD trained networks, across all levels of pruning (Fig. 2B). To understand why, we examined the cosine similarity between the original and the pruned solutions, as this measure has been previously shown to correlate with robustness to pruning in artificial neural networks (Mason-Williams and Dahlqvist, 2024), and, as a measure of weight similarity, it also gives an insight into how much a network has to be re-trained to achieve pre-pruning accuracy. Indeed, we found that the EG trained networks had a higher pruned-unpruned cosine similarity compared to the GD trained networks, in-line with their better post-pruning accuracy (Fig. 2C). Thus, learning with EG produces networks that are more robust to synaptic pruning.

Of course, biological synaptic pruning occurs over the course of development and animals continue learning with sparsified networks. Therefore we next asked whether there were differences between EG and GD when relearning the tasks after pruning. To test this, we took the RNNs trained on the cognitive tasks that had 92.5%, 95% and 97.5% of their synapses pruned and trained them for 1,000 more updates (see Methods for details). We observed that the EG networks were better at relearning the tasks, achieving higher accuracy more rapidly (Fig. 2D-F, top). As well, the EG network weights maintained a higher cosine similarity with the initial pruned weights during relearning (Fig. 2D-F, bottom), indicating that the EG networks had to change their solutions less to reach the new optimal weight-space solutions to the tasks after pruning. Altogether, these results demonstrate that EG is better for maintaining performance and relearning after synaptic pruning. Moreover, they show that in addition to providing a better phenomenological match to the brain, there are biologically relevant learning situations in which EG is better than GD.

### EG is better at ignoring irrelevant features

The previous synaptic pruning experiments showed that EG finds solutions that rely on fewer synaptic weights than the solutions found by GD. Therefore, we hypothesised that finding solutions with a few large weights may also be advantageous for quickly learning tasks where only a few inputs are relevant. Furthermore, EG and its precursors have been shown to have better convergence guarantees in similar optimisation settings (Kivinen and Warmuth, 1997; Littlestone, 1988). Such tasks, involving sparsely relevant information, are highly pertinent for neural circuits as neurons receive both high levels of background noise and inputs from multiple brain areas, not all of which will be always relevant in all contexts (Mante et al., 2013). Furthermore, despite receiving thousands of synaptic connections, generally only a few active synapses are required to drive a cell to fire action potentials (Ikegaya et al., 2013).

To test if EG is beneficial for learning with sparsely relevant synaptic inputs, we first investigated learning in a very simple setting. We modelled a single neuron as a rate-based point neuron receiving either 200, 2000, or 20000 random binary inputs – i.e. presynaptic activations of 1 or 0. We trained the neuron to be active whenever more than 50 of 100 randomly designated relevant synaptic inputs were active (Fig. 3A). For GD, this is equivalent to using the delta rule (Widrow and Hoff, 1988). To solve this task, the neuron needs to learn to ignore irrelevant synaptic inputs and assign large weights only to the relevant inputs. We found that when the total number of synaptic inputs was similar to the number of relevant inputs, EG and GD learning trajectories were also similar (Fig. 3B left). However, as the ratio of relevant inputs to irrelevant inputs decreased, gradient descent struggled to learn which of the inputs were relevant, whereas EG learned much more efficiently (Fig. 3B middle, right).

**Figure 3:**
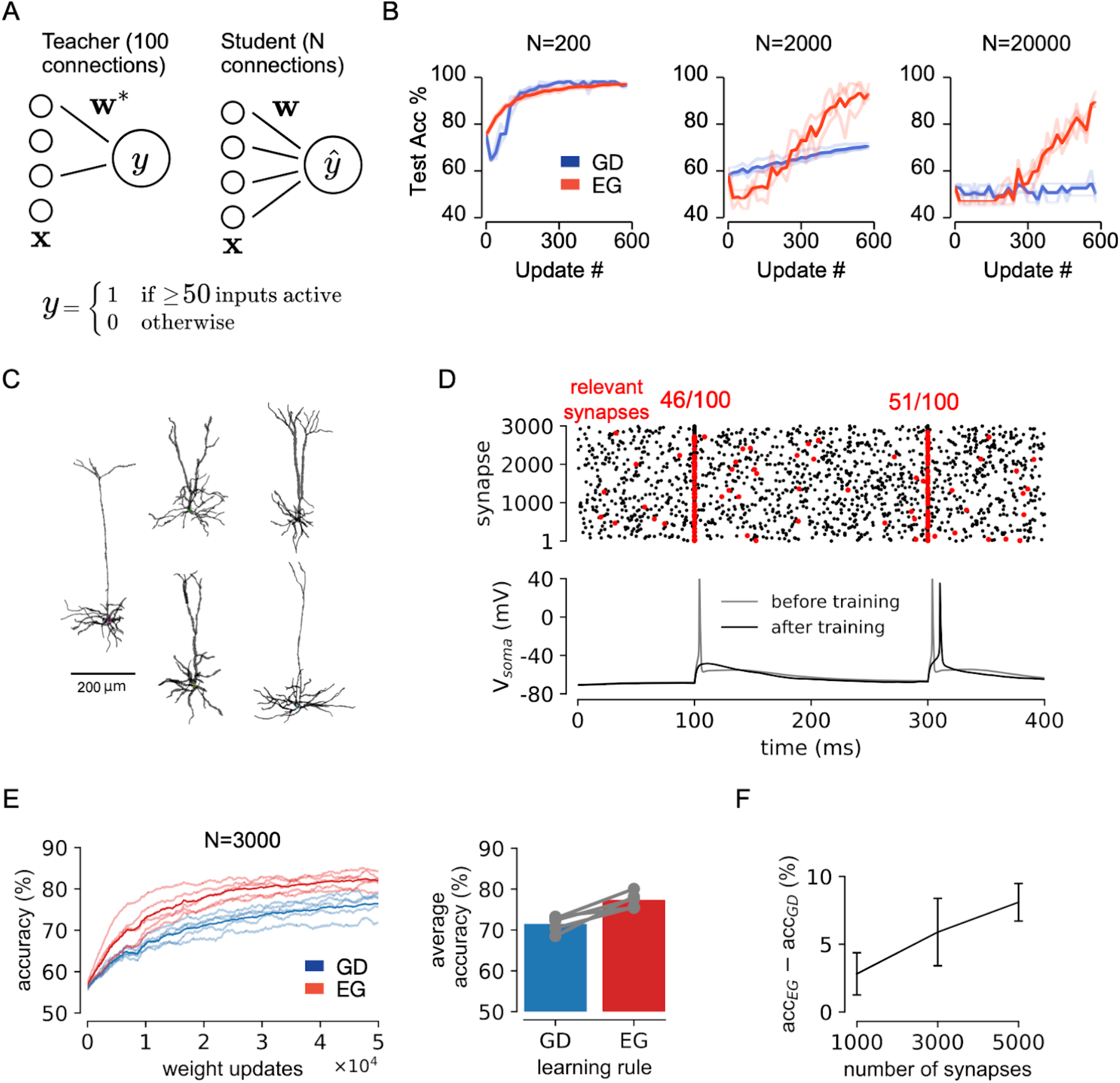
Exponentiated gradient (EG) outperforms gradient descent (GD) when relevant inputs are sparse. **a**| Task description: student neurons receive Bernoulli-distributed binary inputs with N dimensions, and are required to match the output of teacher neurons with 100 non-zero weights. Teacher output is 1 if 50 or more binary inputs are active and 0 otherwise. **b**| Student neuron performance as N increases. n=3 **c**| Morphologies of the biophysically detailed multi-compartment mouse pyramidal neuron models used in experiments. **d**| Example simulation showing a model neuron’s response to two impulse stimuli before and after training. **e**| Left: running average error for neurons trained with EG or GD for N=3000 synapses. Right: accuracy for individual models trained with EG or GD, averaged over the entire task. **f**| Difference in average accuracy increases with the total number of synapses. Error bars are s.d. from n=5 models. (EG orange, GD blue)

While striking, these point neuron experiments (Fig. 3A,B) abstract away a vast amount of biological detail and complexity. Hence, we next tested whether the learning properties of EG persisted in more realistic, spiking, multi-compartment models of pyramidal neurons with conductance-based synaptic inputs distributed across the dendritic tree (Fig. 3C). Again, the model neurons were required to learn to activate only when more than 50 of 100 relevant inputs were active and to ignore irrelevant inputs (Fig. 3D). In this case, optimisation is far more complicated, due to the fact that gradients have to be propagated throughout the entire multi-compartment structure (Bicknell and Häusser, 2021). Nonetheless, we again found that learning with EG was advantageous relative to GD across five different neuron morphologies (Fig. 4E). Furthermore, similar to our point neuron experiments, again we found that as the ratio of relevant inputs to irrelevant inputs decreased, the benefits of learning with EG relative to GD also grew (Fig. 4F). Together, these results demonstrate that learning with EG is beneficial for neurons learning from a sparse set of relevant inputs.

**Figure 4:**
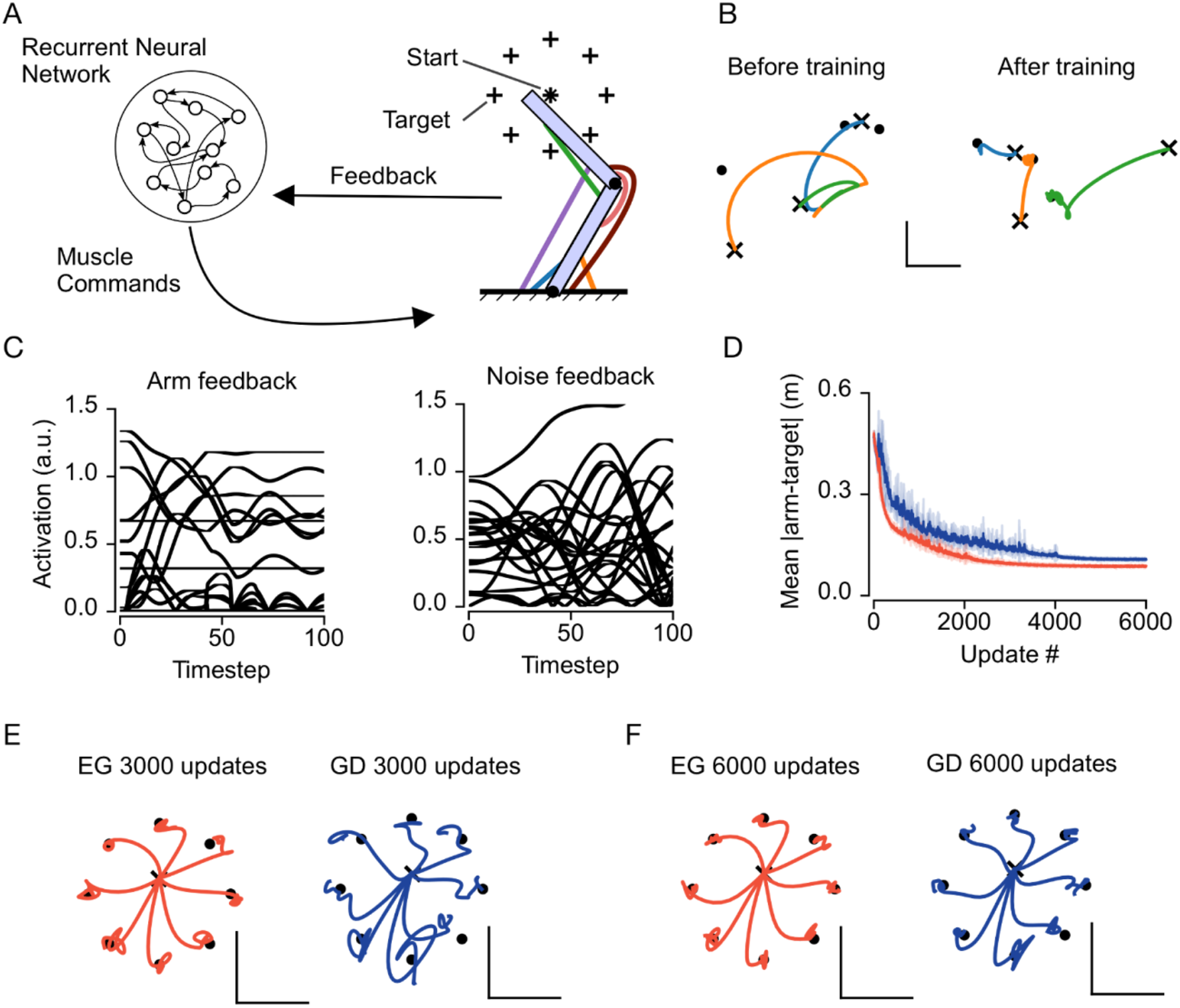
Continuous control with noise is better with exponentiated gradient (EG). **a**| Schematic of an RNN controlling a two-joint planar arm through 8 muscles. The network receives delayed visual and proprioceptive feedback (see Methods) **b**| Three example arm reaches from random starting locations (x symbol) to target locations (filled circles) at the start and end of training. Different colours represent different reaches. Scale bars represent 20cm. **c**| Example feedback to the RNN controller. Left: from arm. Right: noise. **d**| Learning curves for RNNs trained with EG or GD. (From n=9 models). **e**| Validation reaches for EG and GD networks after 3 thousand updates. Scale bars represent 20 cm. **f**| As **e** but for 6 thousand updates.

### EG is better at continuous control with irrelevant features

The above multi-compartment neuron model simulations show that EG’s advantage over GD extends through to realistic levels of biological complexity, albeit for single neurons performing simple tasks. Next, we turned our attention to neural circuits performing more complicated, ethologically relevant tasks in the presence of irrelevant noise. Specifically we considered the task of controlling a two-joint planar arm to perform reaches between random starting and target positions (Fig. 4A, B). This introduces biological complexity to the problem, including a non-linear mapping between the network’s output and the environment, and learning how to employ delayed feedback from the environment to the neural network. As is the case in real brains, we assume that there are many inputs to the circuit that are not relevant to the reaching task, and thus can be treated as noise.

We modelled motor circuits as an RNN receiving a constant stream of visual and proprioceptive inputs from the environment. The output of the RNN was a driving signal to six simulated muscles actuating the arm. In addition to the task relevant visual and proprioceptive signals the network also received additional irrelevant inputs with similar statistics to the task relevant inputs (Fig. 4C). Hence, in order to control the arm accurately, the network must learn to ignore the irrelevant noise.

We compared networks trained with GD and EG in this task and found that there was a clear difference in terms of the speed of learning. Specifically, we observed that EG trained networks (Fig. 3D, orange lines) learned to control the arm faster than GD trained networks (Fig. 3D, blue lines), and converged to a better final solution. EG’s better performance could also be seen by examining a circle of evaluation-reach trajectories after 3,000 and 6,000 updates. We found that EG trained arm controllers showed smooth, direct trajectories to the targets after 3,000 updates, whereas GD trained networks were noticeably less precise (Fig. 3E). Notably, when there were no irrelevant noise inputs the two algorithms were equivalent in their speed of learning (Fig. S4). Hence, EG leads to faster and better learning in the presence of irrelevant inputs, even when applied to non-trivial sensorimotor control tasks.

## Discussion

In this paper we proposed that exponentiated gradient (EG) provides a better algorithm for training computational neuroscience models than gradient descent (GD). We presented three lines of evidence justifying this claim. First, we highlighted the theoretical principles shared by both learning rules, and empirically showed that EG is no less powerful than GD at learning a challenging battery of cognitive tasks (Fig. 1). Second, we found that EG matches neural phenomena better than GD: EG maintains Dale’s Law and log-normal weights, and produces multiplicative, proportional synaptic changes. Third, we observed several advantages of EG over standard GD in biologically relevant task settings, including: a greater ability to adapt to synaptic pruning (Fig. 2), the ability for single neurons to learn when very few inputs are relevant to the task (Fig. 3), and the ability for recurrent networks to learn rapidly when engaged in sensorimotor control in the presence of noise signals (Fig. 4). Altogether, these results support our assertion that EG is a better algorithm than GD for training neural network models of the brain.

### Theoretical considerations

We observed that EG leads to different solutions than GD. What is the cause of these differences? One observation is that using different synaptic proximity penalties (see Section 2.1) effectively alters the “distance” between distinct settings of synaptic weights (Gunasekar et al., 2018). As a result, given the same initial weights, GD and EG have different “distances to travel” to a target weight setting, and as such, in general they may be more inclined to discover some weight configurations over others. That observation aligns with our results in the Mod-Cog suite of tasks (Fig. 1) where we found that the behaviour of the networks was very similar, but the actual synaptic weights learned were quite distinct. Another observation to explain the differences between the algorithms is that the exponential operation in EG leads to more emphasis being placed on certain directions in the gradient, and this effect will likely be compounded by the multiplicative nature of the weight updates (Schwarz et al., 2021). Indeed, this nonlinear credit assignment process will bias the network towards weights where a small number of connections become very strong, which has implications for pruning (Fig. 2) and learning tasks with a small number of relevant inputs (Fig. 3, 4).

These considerations also have interesting implications for questions of network development and initialisation. Recent deep learning experiments have shown that at initialisation networks contain task-specific subcircuits that both change a lot during learning, and are capable of solving the task by themselves (Frankle and Carbin, 2018). As such, it may be that EG is better at discovering these so-called “winning lottery tickets” within a network’s initialisation, i.e. finding the subset of synaptic weights that should change a lot in order to learn the task. Relatedly, one potential disadvantage for EG is that, due to its inability to change the sign of synaptic weights, if the network is initialised in a state where the only feasible solutions require a sign change in some synapses, then GD will outperform EG. We did not observe this in our simulations, though, likely because this situation is only relevant in an under-parameterised regime where the likelihood of having randomly initialised a group of synapses to the right sign for solving the task is much smaller. However there are inhibitory-excitatory networks architectures that learn well in such regimes (Cornford et al., 2021; Haber and Schneidman, 2022), and biological circuits are not under-parameterised.

### Limitations and future work

Our analysis and experiments rely on gradient-based learning signals. While there is ongoing theoretical and experimental work investigating how brain circuits might estimate quantities correlated with the gradient (as they *must* in order to improve on tasks with small synaptic changes (Richards and Kording, 2023)), it is not yet clear how real neural circuits can engage in gradient-like calculations (Lillicrap et al., 2020). However, we emphasise that our central contribution is agnostic to the specific form of the neuronal credit signal – anything that is correlated with the gradient (e.g. reward signals) could be used in place of the explicit gradient. Instead, we leveraged the often implicit concept of synaptic distance, which dictates how such credit signals are used to update synaptic strengths. Nonetheless, future work could examine whether other, more biologically plausible credit assignment mechanisms would lead to the same set of results we obtained here.

Another potential limitation is the difficulty of the tasks. While these tasks are relevant to computational neuroscience (Khona et al., 2023; Yang et al., 2019), they are simple from an optimisation perspective: such tasks are far less complicated than controlling embodied agents in open-ended environments or engaging in real-world language modelling (Brown et al., 2020; Todorov et al., 2012). Previous work has shown that biologically plausible learning algorithms can fail to scale to more difficult tasks (Bartunov et al., 2018), raising the possibility that our conclusions may also change with an increase in task difficulty. However, because GD is known to perform well in more complicated tasks, and EG and similar algorithms have also been shown to scale-up to hard computer vision tasks (Bernstein et al., 2020; Schwarz et al., 2021; Sun et al., 2022), it is unlikely that such a concern applies here. But, we hope future work will explore more fully the representational differences between EG and GD in tasks that are of a real-world level of complexity.

Finally, we did not consider the role of different cell types. Instead we modelled the heterogeneous biological complexity of neural circuits abstractly using a homogenous population of rate-based rectified linear units. Although it is a common practice to make such simplifications in Neuro-AI studies, the question of cell types may be particularly important when considering which synaptic proximity penalty, and therefore learning algorithm, to use. In particular it may be that synaptic plasticity in inhibitory and excitatory neurons should be modelled more accurately using different distance functions, and in line with this, recent work has shown that synaptic dynamics are different for PV+ interneurons and pyramidal neurons (Melander et al., 2021). The biophysical modelling approach we employed for the single-neuron experiments (Fig. 4) is ideal for exploring such cell-type differences in future work.

### Importance for understanding learning in real brains

The use of EG as an algorithm for training models of neural circuits has both computational and experimental implications. First, it could facilitate downstream computational neuroscience studies investigating the functional properties of skewed, log-normally-distributed weights. While there are older works on log-normal-consistent learning (Van Rossum et al., 2000), these studies used synaptic plasticity rules that are unable to learn even the sort of simplified tasks we studied here. Our work provides a means for computational researchers to explore learning with log-normal distributions in weights while maintaining the ability to learn more demanding, or cognitively relevant tasks. Second, this work provides a normative, computational explanation for some previously observed experimental phenomena (e.g. multiplicative weight updates). Finally, we believe that the EG framework, and more generally mirror descent, is an important step towards data-constrained learning algorithms. Here, it allowed us to incorporate several biological observations without sacrificing the performance of GD. In future work, we hope this framework will co-evolve with experimental observations to develop better models of learning in neural circuits.

## Acknowledgements

We would like to thank Daniel Levenstein, Colin Bredenberg, Zahid Padamsey, Leonid Savtchenko, and Dongyan Lin for their helpful comments on the manuscript and discussion. We would also like to thank all members of the Richards and Lajoie labs for creating a supportive and intellectually stimulating environment.

## Funding Acknowledgement

This work was supported by NSERC (Discovery Grant: RGPIN-2020-05105; Discovery Accelerator Supplement: RGPAS-2020-00031; Arthur B. McDonald Fellowship: 566355-2022); CIFAR (Canada AI Chair; Learning in Machine and Brains Fellowship) and IVADO (IVADO postdoctoral fellowship). This research was enabled in part by support provided by (Calcul Québec) (https://www.calculquebec.ca/en/) and the Digital Research Alliance of Canada (https://alliancecan.ca/en). The authors acknowledge the material support of NVIDIA in the form of computational resources. The authors acknowledge the use of the UCL Myriad High Performance Computing Facility (Myriad@UCL), and associated support services, in the completion of this work.

## Method details

All simulations were implemented and results analysed using Python. Neural networks were implemented in PyTorch, and multi-compartment neuron models in NEURON. Full code, model checkpoints, hyperparameter settings, and training data will be released upon publication. Simulations were run on either MILA Quebec AI institute’s compute cluster, or University College London’s Myriad high performance computing system. The majority of ANN experiments were run using either NVIDIA RTX8000 or A100 GPUs.

### Recurrent Neural Network Architecture

For Mod-Cog (Sec. 2.2) and continuous control (Sec. 2.4) experiments, we modelled all recurrent neural networks (RNNs) as follows. At each time t and input *x*^*t*^, the hidden state *h*^*t*^ of 2.5 thousand neurons was defined as:

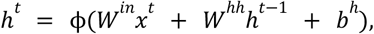

where ϕ is the neuronal nonlinear activation function. Throughout we set ϕ to be the rectified linear function (ReLU), in order to constrain activity to be positive only and to capture the rectification operation performed by biological neurons in low activity regimes. The values of *x*^*t*^ were task dependent and at time t=0 the hidden state was set to be a vector of zeros.

The output activation of the RNN at each time point is:

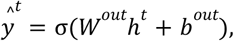

where the choice of σ and the dimensionality of *W*^*out*^ is dependent on the task (see task details below).

All network parameters (*W*^*out*^ ∈ *R*^|*h*| ×|*h*|^, *W*^*hh*^ ∈ *R*^|*h*| ×|*h*|^, *W* ^*in*^ ∈ *R*^|*x*| ×|*h*|^, *b*^*out*^ ∈ *R*^|*y*|^, *b*^*h*^ ∈ *R* ^|*h*|^, where |*h*| denotes the cardinality of *h*) were randomly initialised using the same distribution for EG and GD. For *all* weights, we initialised each individual weight to be log-normal with

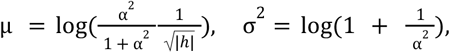

where α = 1. 5. The resulting weights were positive with mean 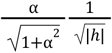 and variance 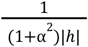, so to obtain final weights we randomly multiplied them by ± 1 with equal probability. This makes the weights zero mean, and due to the variance of *XY* for independent *X* (positive weights) and *Y* (signs), the variance becomes 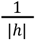 (the same for input, hidden, and output weights). For any positive α, this initialization scheme produces the same mean and variance of the final weights distribution. We set α = 1. 5 to roughly reproduce the shape of experimentally observed log-normal synaptic weights and did not observe that network training was sensitive to this choice (not shown).

For both biases, for GD, we initialised them from a uniform distribution 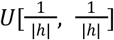 (standard PyTorch initialisation). In order to account for the inability of EG to change parameter sign, models optimised by EG we initialised to have two bias vectors, one positive, one negative: where we initialized 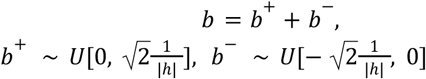. For both GD and EG, the biases therefore were mean zero with 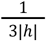 variance.

### Learning algorithm and optimisation details

For all experiments, apart from the multi-compartment neuron models, parameters were updated using gradients calculated using PyTorch’s automatic differentiation engine, autograd. For RNNs gradients were calculated using backpropagation through time (BPTT) with no truncation. Parameter updates were then obtained according to either gradient descent (standard BPTT) or exponentiated gradient as described in the main text. We applied momentum *α* (in the standard PyTorch way for SGD) and weight decay *γ*:

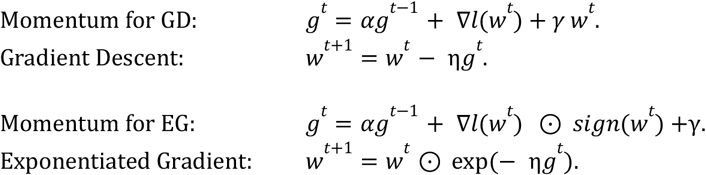

For EG, the sign of weights ensures both negative and positive weights change correctly (a positive gradient ∇*l*(*w*^*t*^) means the weight should decrease, and without the sign *w*^*t*^ ⊙ exp(− η∇*l*(*w*^*t*^)) would instead increase the negative weights). We further discuss the derivation of generic mirror descent and the weight decay we use for EG in the Supplementary.

### Training details

For all experiments we stabilised network training by constraining the norm of the gradient vector to be less than or equal to a scalar value that was chosen via hyperparameter search (Pascanu et al., 2012). Training hyperparameters, such as learning rate, momentum, etc. were selected using grid search to minimise the final validation loss. Please see the individual task descriptions for specifics and further details.

### Task-specific details

#### Mod-Cog

All 82 tasks comprising the Mod-Cog dataset of cognitive tasks shared the following input and output structure (see S1 for task examples). Network inputs were a concatenation of 4 vectors: fixation, stimulus modality 1, stimulus modality 2, and rule encoding. The fixation input was a one dimensional variable that remained at 1 for the duration of a trial until the onset of the decision (response) period when it dropped to 0. The two input stimulus modalities were both ∈ *R*^16^ and each encoded a one dimensional circular variable with preferred directions uniformly spaced from 0 to 2π. Input stimuli were presented on one or both input rings for trial time-periods in a task dependent manner as detailed below. Finally, throughout each trial, the rule encoding was a constant one-hot ∈ {0,1}^84 vector indicating the current task to be performed.

For Mod-Cog network outputs, *ŷ*, were ∈ *R*^17^, comprising a scalar fixation output and an output ring ∈ *R*^16^. During the response period, output ring unit activities were converted to probabilities using a softmax function and trained using a cross entropy loss to match a one-hot target vector *y*, representing the location of target activity on the output ring. Before the response period, network outputs were trained to match the target vector *y* using a mean squared error loss.

To maximise computational efficiency task trials were drawn from neurogym objects (Molano-Mazon et al., 2022) and stored as a dataset as follows. Trials were drawn randomly from one of the 82 tasks, and concatenated together until a sequence length of 350 was obtained. 512 of these sequences were then stacked to form a batch. The dataset itself was composed of 1000 of these batches. Therefore 1000 updates corresponds to one pass through the dataset. Validation and test datasets of 100 batches were additionally drawn.

We trained networks for 6000 updates (i.e. 6 passes through the dataset). During training the learning rate followed a warmup (from 0), constant, cooldown schedule (to 0). Warmup and decay periods were 500, 1000 updates respectively. Best hyperparameters were selected based on average final loss over three random seeds calculated on the validation dataset. These settings were then used for five more random seeds, and final results were reported on the held out test dataset. During training gaussian noise (mean 0, std deviation 0.1) was added to the task inputs. Validation and test sets were evaluated without noise. The subtask specific details are postponed to the Supplementary.

#### Mod-Cog Pruning

For all pruning experiments non-bias network parameters (i.e. weights) were pruned according to L1 global unstructured pruning. As such, for a given prune percentage *x*, all non-bias parameters were sorted according to their absolute value, and the smallest *x* percentage of parameters were masked and set to 0. Networks were pruned after 5 thousand updates. For retraining experiments, the networks continued to learn with the pruning mask in place (i.e. pruned connections were held at 0) for the final thousand updates. For further details see the documentation for “torch.nn.utils.prune”.

#### Sparsely relevant inputs

##### Point neuron models

A student neuron received N independent Bernoulli inputs (p=0.5). Out of the N inputs, 100 were randomly designated to be relevant by a teacher neuron, whose weights were 0 for irrelevant inputs, and 1 for relevant inputs. For each activation, the output label from the teacher neuron was 1 if at least R=50 of those inputs were active.

Student neuron weights were initialised to be *r*/(*p* * *n*). For the student neuron, its output *x*^*⊤*^ *w* was transformed through a sigmoid activation function as

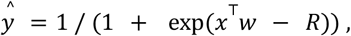

and then passed to the cross-entropy loss. Therefore, for *x*^⊤^ *w* that exceeded R the *ŷ* was larger than 0.5, predicting *y* = 1.

For each N, we constructed a training dataset of 10,000 inputs, and validation, test datasets of 1000 inputs. Neurons were trained for 600 iterations with a batch size of 500 and hyperparameters chosen through grid search and best final accuracy on the validation set. Results were reported over 3 seeds.

##### Multi-compartment neuron models

We used six biophysically detailed models of mouse pyramidal neurons from the Allen Cell Types Database IDs: 483061182, 483101699, 486560376, 484564503, 485836906 and 468193142] (Allen Cell Types Database, 2015). Each model comprises a 3-dimensional morphology reconstructed from experiments and is endowed with a suite of 10 Hodgkin-Huxley style ion channel conductances at the soma. The parameters of each model have been fitted to patch clamp recordings associated with its unique morphology; for full details, see (Gouwens et al., 2018).

Excitatory synapses with AMPA and voltage-dependent NMDA conductances, and inhibitory synapses with GABA conductances were distributed uniformly across the basal dendrites. Synapse models were as described in (Bicknell and Häusser, 2021). For a pre-synaptic spike arriving at time *t*_0_, and local dendritic membrane potential *V*_*loc*_ (*t*), the synaptic current from a given excitatory or inhibitory synapse is described by

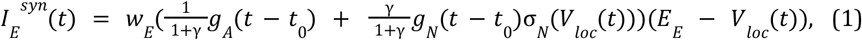

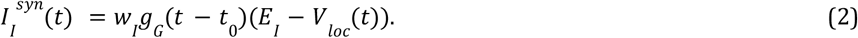

In Eqs. (1-2), *w*_*E*_ and *w*_*I*_ denote synaptic weights, *γ* is the NMDA/AMPA ratio, *E*_*E*_ and *E*_*I*_ denote reversal potentials, and the functions *g*_*A*_, *g*_*N*_, *g*_*G*_ describe double-exponential activation kinetics of the AMPA, NMDA and GABA conductances,

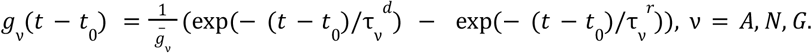

The normalizer 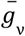 ensures that the activation peaks at a magnitude of 1. The voltage-dependence of the NMDA conductance, σ_*N*_ (*V*_*loc*_), is modelled by the sigmoid

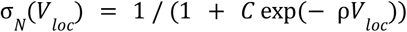

using parameters *C* = 1/3. 57, ρ = 0. 062, as described by (Jahr and Stevens, 1990).

We set γ = 2 for the NMDA/AMPA ratio, and *E* _*E*_= 0 *mV*, and *E*_*I*_ =− 75 *mV* for the reversal potentials. The rise and decay time constants of the activation kinetics were as follows. AMPA: 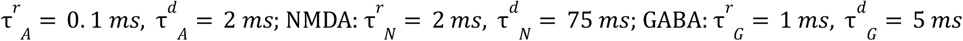.

###### Task

We consider *N* total synapses, each receiving independent Poisson spike trains at spontaneous rates of 1.25 spikes/s. A number K of the excitatory synapses are designated as “relevant”, to which the neuron should pay attention, whereas the remaining synapses are a source of noise. After a 100 ms burn-in period, an impulse stimulus is presented, with each synapse randomly receiving an input spike with probability *p*_*stim*_. The classification task requires the neuron to fire an action potential whenever at least *R* out of the *K* relevant synapses are activated at stimulus time. We fixed the parameters *p*_*stim*_ = 0. 5, *R* = 50, *K* = 100, while varying *N*.

The task was run for 60,000 total stimulus presentations with learning as described in the next section. Performance over the training period was quantified as the fraction of correct classifications in blocks of 40 presentations. For one of the model neurons (ID: 483061182) learning did not converge for either the GD or EG algorithm; this model was excluded from further analysis.

###### Learning rules

Learning was implemented using the gradient-based technique of (Bicknell and Häusser, 2021), generalised for use with the Allen Institute models.

In the binary classification task described above, a neuron is required to spike for one class of input patterns, which we denote ⊕, and remain silent for the other, denoted ⊖. Learning proceeds according to the Tempotron principle (Gütig and Sompolinsky, 2006): When a classification error occurs, synaptic weights are updated with the aim of pushing the somatic membrane potential, *V*_*soma*_, above spike threshold for ⊕ patterns, and below threshold for ⊖ patterns. To do this efficiently, the updates must depend on the derivative of *V*_*soma*_ with respect to the weights, ∂*V*_*soma*_ /∂*w*_*i*_. In this framework, the GD and EG learning rules therefore take the forms

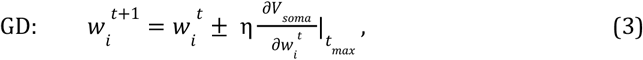

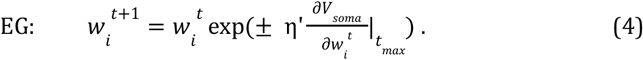

In Eqs. (3-4), the symbol ± denotes that the sign is positive for ⊕ patterns and negative for ⊖ patterns, η and η’ are learning rates, and the derivative is evaluated at the time of maximum membrane potential during the pattern presentation, *t*_*max*_. For ⊕ patterns, *t*_*max*_ is the time at which the subthreshold potential reaches its peak value, whereas for ⊖ patterns, *t*_*max*_ is the time of the first action potential (in practice, 2 ms before the action potential, for numerical stability). Note that these updates are only performed if there has been an error on presentation *t*; weights are left unchanged after successful classifications.

Computing ∂*V*_*soma*_ /∂*w*_*i*_ for the biophysical models is nontrivial, as the derivative must be propagated through a system of differential equations that governs the membrane potential dynamics. In general, suppressing the technical model details, each neuron is described by a dynamical system

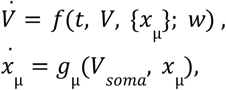

where the vector *V* denotes the membrane potential in each of the many compartments of the discretized morphology, the variables *x*_µ_ are Hodgkin-Huxley gating variables for the somatic ion channels (µ_µ_ = 1, 2, …; typically two gating variables per channel species), *f, g*_µ_ are nonlinear functions describing current flow and ion channel kinetics, and *w* is a vector of synaptic weights. The compartments of the morphology are ordered such that the somatic membrane potential is given by the first element of *V*, i.e. *V*_*soma*_ ≡ *V*_1_.

Taking a derivative of the equation above with respect to a weight, *w*_*i*_, applying the chain rule, and swapping the order of the time and weight derivatives, leads to the system

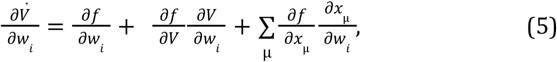

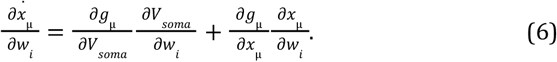

Eqs. (5-6) describe how the membrane potential in each compartment depends on *w*_*i*_ over time. By solving it numerically for each weight, in parallel with simulations of the model, the crucial gradient terms can be read off as the first component of ∂*V*_*soma*_ /∂*w*_*i*_ (*t*_*max*_).

For efficient application in the classification task, we simulated the models using the NEURON simulation environment (Hines and Carnevale, 1984), while solving the last equation for the gradients using a custom-written Python package. Learning rates η and η’ were selected via grid search to maximise task performance averaged over the entire training period.

#### Continuous Control

For continuous control, all trials were randomly generated and therefore hyperparameters were chosen based on grid search performance and experiments with 5 seeds re-run for final results.

For the continuous control experiments, the RNN output layer consisted of six neurons with a sigmoid nonlinearity to ensure positive outputs. The output from this last layer served as a muscle drive that was passed to the six muscle actuators controlling the bi-planar arm. The environment models employed were from the open-source Python package motornet (Codol et al., 2024). Briefly, the actuators were six Hill-type muscle models, which produced forces based on their previous activation and the incoming muscle drive from the network. These forces were applied to the effector’s bones proportionally to their insertion points onto it. The new bone positions were then used to determine endpoint positions in the Cartesian space.

Specifically, we employed the *RigidTendonArm* effector with default parameter values and the Euler algorithm for numerical integration. The muscle actuators were *MujocoHillMuscle* objects with passive force contribution set to zero. The simulation time constant was set to 10 ms for all models.

The RNN received as input a vector containing the Cartesian coordinates of the arm’s endpoint, start position and target position, as well as each actuator’s length and velocity. During training the visual feedback (the arm’s current endpoint) was delayed by 50 ms (5 simulation steps) and the proprioceptive feedback (each actuator’s length and velocity) was delayed by 20 ms (2 simulation steps) to match feedback delays observed in the motor cortex of biological organisms (Omrani et al., 2016).

Networks were trained to minimise a composite loss quantifying performance over each 1 second reach:

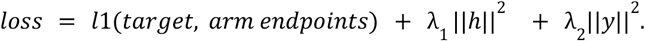

Here *l*1(*target, arm endpoints*) is the mean of the l1 distance between the arm endpoint and target location over the 1 second reach. λ_1_ and λ_2_ penalise large RNN hidden states and muscle drives respectively, and were both set to 0.1. These environment states are used to compute the gradient as the environment built through MotorNet is differentiable. This is equivalent to backpropagating through an accurate forward model of the effector.

To model learning in the presence of irrelevant inputs, the RNN received 500 time-varying noise inputs *x*_*t*_ (at time *t*) in addition to the arm feedback. These noise inputs obeyed the following coordinate-wise dynamics:

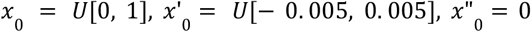

For *t* > 0, if *x*_*t*_ = 0, then we set the “speed” and “momentum” 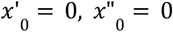. After that, for *θ* = 0. 0005,

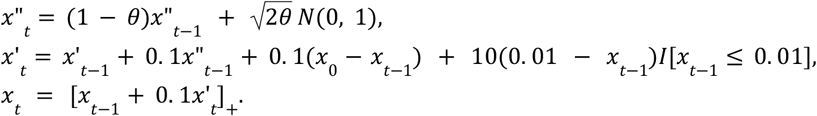

Essentially, the noise inputs *x*_*t*_ are sampled by sampling the noise “momentum” from an Ornstein–Uhlenbeck process, the noise “speed” integrates the momentum variable and additionally pushes *x*_*t*_ away from zero and towards the initial values, with *x*_*t*_ itself integrating the speed variable. If *x* reaches zero, its speed and momentum are reset.

### Analysis details

To compare task accuracies (Fig. 1) and weights during pruning (Fig. 2), we used the cosine similarity measure between two vectors *x, y*,

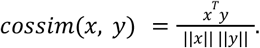

For comparing weight distributions (Fig. S3), we used the one-sample Kolmogorov-Smirnov test statistic (Hollander et al., 2015) to compare distributions with a normal distribution,

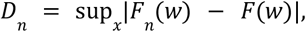

where *F* (*x*) is the CDF of the (z-scored, i.e. normalised by the sample mean and standard deviation) weights and *F*(*x*) is the CDF of the standard normal.

For comparing two pairs of weights distribution (Fig. S3), we used the two-sample Kolmogorov-Smirnov test statistic (Hollander et al., 2015) for CDFs of z-scored weights *F*_*n*_ and *G*_*n*_,

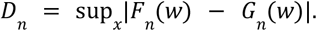

For fitting the slope of eigenvalues (Fig. S2B), we used code made available by (Agrawal et al., 2022).

## Supplementary

## Appendix A: Mirror descent

In this section we provide a more technical overview of mirror descent and weight decay.

### Mirror descent derivation

In this section, we provide an introduction to mirror descent. Several textbooks, such as (Bubeck, 2015; Shalev-Shwartz and others, 2012), provide a more detailed discussion on this framework.

In standard gradient descent, to minimise a loss function l(w), a single update starting from weight *w*^*t*^ is derived as follows:

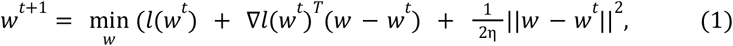

where η is the learning rule. Therefore, a single step of GD linearises the loss at the current weight *w*^*t*^, and then adds a quadratic term to the linearisation to establish a region in which linearisation is considered accurate (which is controlled by η, which can be viewed as an alternative interpretation of the learning rate).

The problem in (1) is convex, and we can find its solution analytically, getting a gradient descent step,

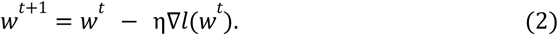

In mirror descent (introduced in (Nemirovskij and Yudin, 1983)), the weight update is regularised by a different penalty *D* _ϕ_ (*w, w* ^*t*^),

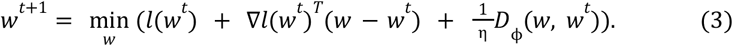

What should *D* _ϕ_ (*w, w* ^*t*^) be to work as a penalty? Like for the two-norm in Eq. 1, we expect it to be nonnegative, zero at *w* = *w*^*t*^, and increase as *w* moves away from *w*^*t*^.

Mirror descent formalises this intuition of a penalty with Bregman divergence (Bregman, 1967). To define it, we first take a differentiable function ϕ(*w*) that is strictly convex, i.e. a function that follows for *w*’ ≠ *w*

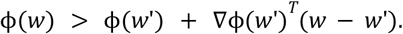

For such functions, we can define the Bregman divergence as the difference between the function and its linearisation:

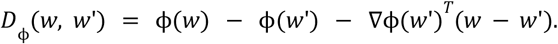

It’s worth noting that mirror descent is usually defined for strongly convex functions (Bubeck, 2015), i.e. the ones following

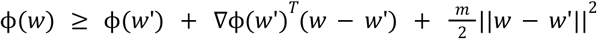

for some constant *m*. This is helpful for deriving algorithmic convergence guarantees (regret bounds). However, to derive the update itself, we only need strict convexity to guarantee invertibility of ∇ϕ. Another way to view this requirement is by examining the minimisation problem in Eq. 3: for a strictly convex potential, the corresponding Bregman divergence is also strictly convex, so adding it to a linear function (which is the linearised loss) makes the overall problem have a unique minimum.

To derive a closed-form mirror descent update, we differentiate the functional in Eq. 3 w.r.t. *w* and equal the result to zero. As it’s strictly convex (as a sum of convex functions and a strictly convex one), this will give us a unique minimum:

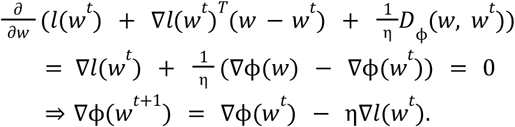

The first example of mirror descent is the regular gradient descent. For 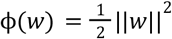, we have that ∇ϕ(*w*) = *w*, and therefore get the gradient descent update in Eq. 2.

To derive exponentiated gradient descent, we need to split the weights into positive and negative ones. One way to do that is to take a vector of signs *s* and a vector of positive weight *w*^+^, such that *w* = *s* ⊙ *w*^+^ (where ⊙ stands for element wise multiplication). We will only optimise over *w*^+^ and hold *s* fixed. We then take ϕ(*w*^+^) = (*w*^+^) log *w*^+^, so Eq. 3 becomes

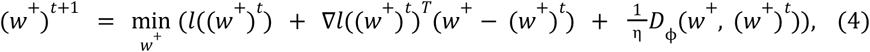

and therefore

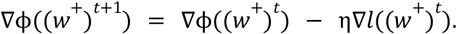

By the chain rule,

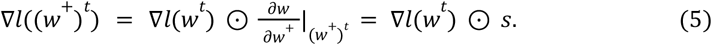

Then, to derive the final update, we note that ∇ϕ(*w*^+^) = 1 + log *w* ^+^, and therefore from Eqs. 4-5 we have that

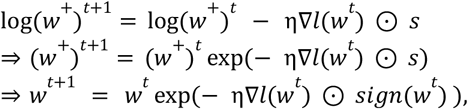

where the last line uses the fact that the weight has a fixed sign.

### Exponentiated gradient with weight decay

For both GD and EG, we added weight decay such that it decreases the current weight *w*^*t*^ as *γ*’*w*^*t*^, where for GD *γ*’=1 − η*γ* and for EG *γ*’=exp(− η*γ*). In terms of the mirror descent formulation, we can view it as

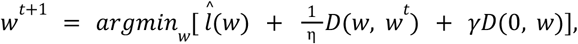

which for GD recovers the standard γ||*w*||^2^ weight penalty.

## Appendix B: Mod-Cog subtask specific details

The set of 82 Mod-Cog tasks were introduced by Khona et. al. 2023. The Mod-Cog dataset extends an original set of 20 cognitive tasks from Yang et. al. 2019 by including sequence generation and interval estimation requirements. The reader is referred to these references for full details including experimental motivation, here we provide a summary. The 20 original tasks can be split into 4 families: Go, Decision Making, Delay Decision Making and Matching. For all tasks, the network is required to match a target output direction for the duration of the response period. Mod-cog adds 40 additional tasks by requiring the network to instead output a sequence (clockwise or anticlockwise) during the response period, with the sequence beginning at the base task’s target output. Furthermore, 11 of the original Yang et al. 2019 tasks involve delays periods. Mod-cog adds a further additional 22 tasks by offsetting the original task’s target output (clockwise or anticlockwise) by a magnitude dependent on the length of the delay period. See Fig. S1 examples.

Here we provide further details regarding the base set of cognitive tasks introduced by Yang et. al. 2019 (text adapted from Yang 2019).

### Go Task family (6 tasks)

[Go, Reaction Time Go, Delay Go, Anti Go, Anti Reaction Time Go, Anti Delay Go]. A single stimulus is presented randomly on either input modality 1 or modality 2, and the target response is in the same direction as the stimulus unless “Anti” when it is the opposite. For non-Reaction Time, the input appears before the fixation cue goes off, whereas for Reaction Time the input appears when the fixation goes off and the network should respond as soon as the stimulus appears. For the delay versions of the task, the input stimulus ends before the fixation drops with a random delay.

#### Decision making family (5 tasks)

[DM 1, DM 2, Ctx DM 1, Ctx DM 2, MultSen DM tasks]. In each trial, two stimuli are shown simultaneously and are presented till the end of the trial. Stimulus 1 is drawn randomly between 0 and 360°, while stimulus 2 is drawn uniformly between 90 and 270° away from stimulus 1. In DM 1, the two stimuli only appear in modality 1, while in DM 2 the two stimuli only appear in modality 2. In DM 1 and DM 2, the correct response should be made to the direction of the stronger stimulus (the stimulus with higher γ). In Ctx DM 1, Ctx DM 2 and MultSen DM tasks, each stimulus appears in both modality 1 and 2. In the Ctx DM 1 task, information from modality 2 should be ignored, and the correct response should be made to the stronger stimulus in modality 1. In the Ctx DM 2 task, information from modality 1 should be ignored. In the MultSen DM task, the correct response should be made to the stimulus that has a stronger combined strength in modalities 1 and 2.

#### Delay decision making family (5 tasks)

[Dly DM 1, Dly DM 2, Ctx Dly DM 1 and Ctx Dly DM 2]. These tasks are similar to the corresponding tasks in the DM family, except that in the Dly DM family tasks, the two stimuli are separated in time. The two stimuli are both shown briefly and are separated by a delay period. Another short delay period follows the offset of the second stimulus.

#### Matching family (4 tasks)

[DMS, DNMS, DMC, DNMC]. In these tasks, two stimuli are presented consecutively and separated by a delay period. Each stimulus can appear in either modality 1 or 2. The network response depends on whether or not the two stimuli are ‘matched’. In the DMS and DNMS tasks, two stimuli are matched if they point toward the same direction, regardless of their modalities. In DMC and DNMC tasks, two stimuli are matched if their directions belong to the same category. The first category ranges from 0 to 180°, while the rest from 180 to 360° belong to the second category. In the DMS and DMC tasks, the network should respond toward the direction of the second stimulus if the two stimuli are matched and maintain fixation otherwise. In the DNMS and DNMC tasks, the network should respond only if the two stimuli are not matched, that is, a non-match, and fixate when it is a match.

## Appendix C: Supplementary Figures

**Supplementary Figure 1:**
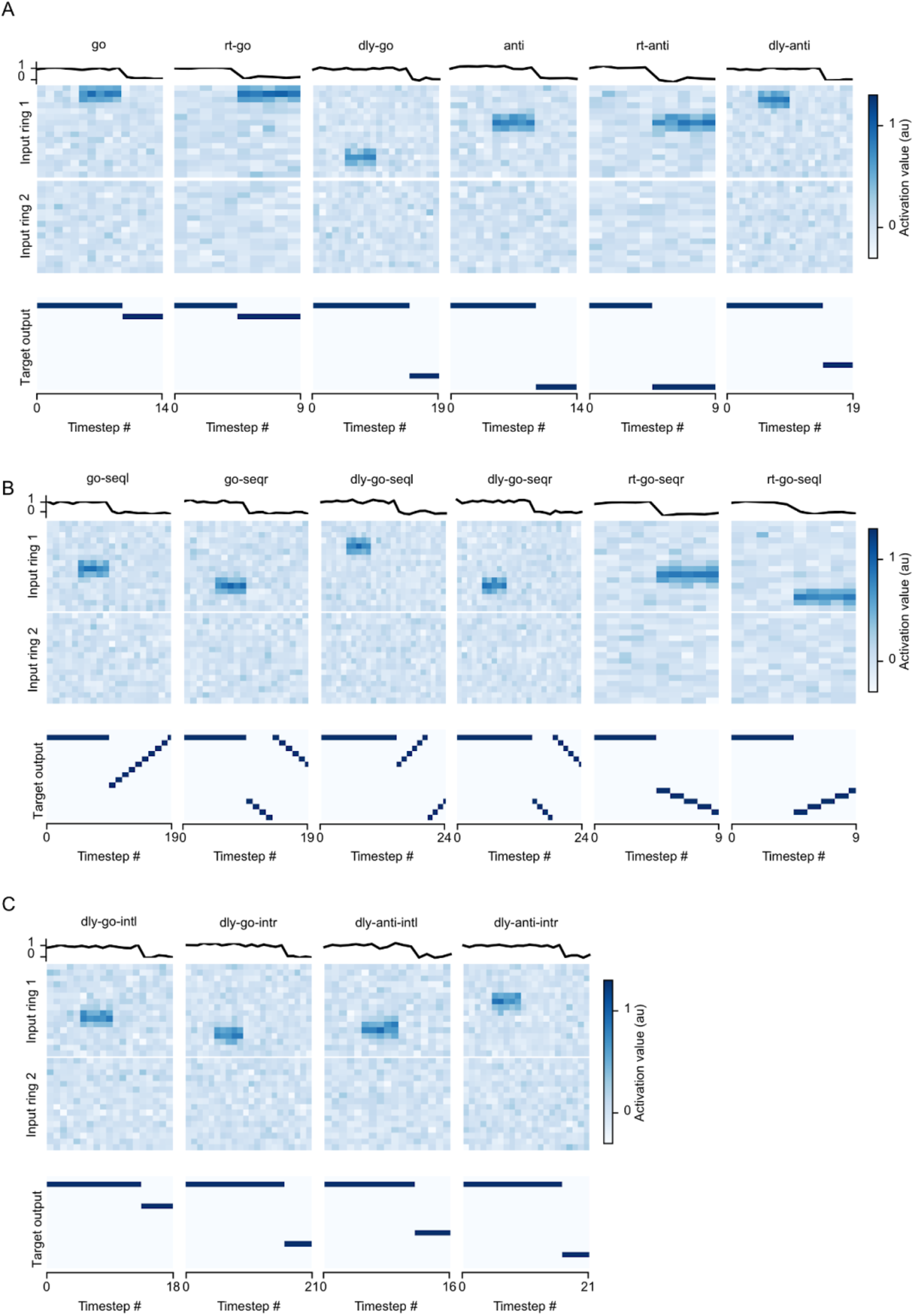
Mod-Cog task examples from Go task. **a**| Go task family. **b**| Mod-cod sequence generation applied to go task family. **c**| Mod-cod delay estimation with the go task family.

**Supplementary Figure 2:**
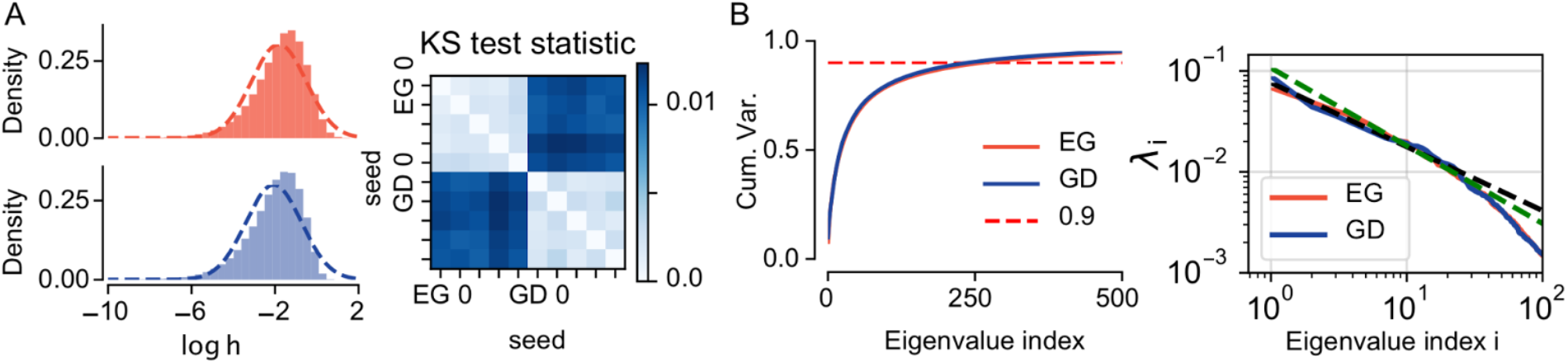
GD and EG trained network activity when solving. **a**| Histograms of EG (red) and GD (blue) trained network activity in log-space. Dashed lines show normal fit, data is a random sample of 1000 time points at the end of training. **b**| Spectral properties of data shown in **a**. (Left) Cumulative explained variance vs eigenvalue. (Right) Log-log plots of eigenvalue index vs value. Dashed lines show linear fit (fit from 3rd to 30th eigenvalues).

**Supplementary Figure 3:**
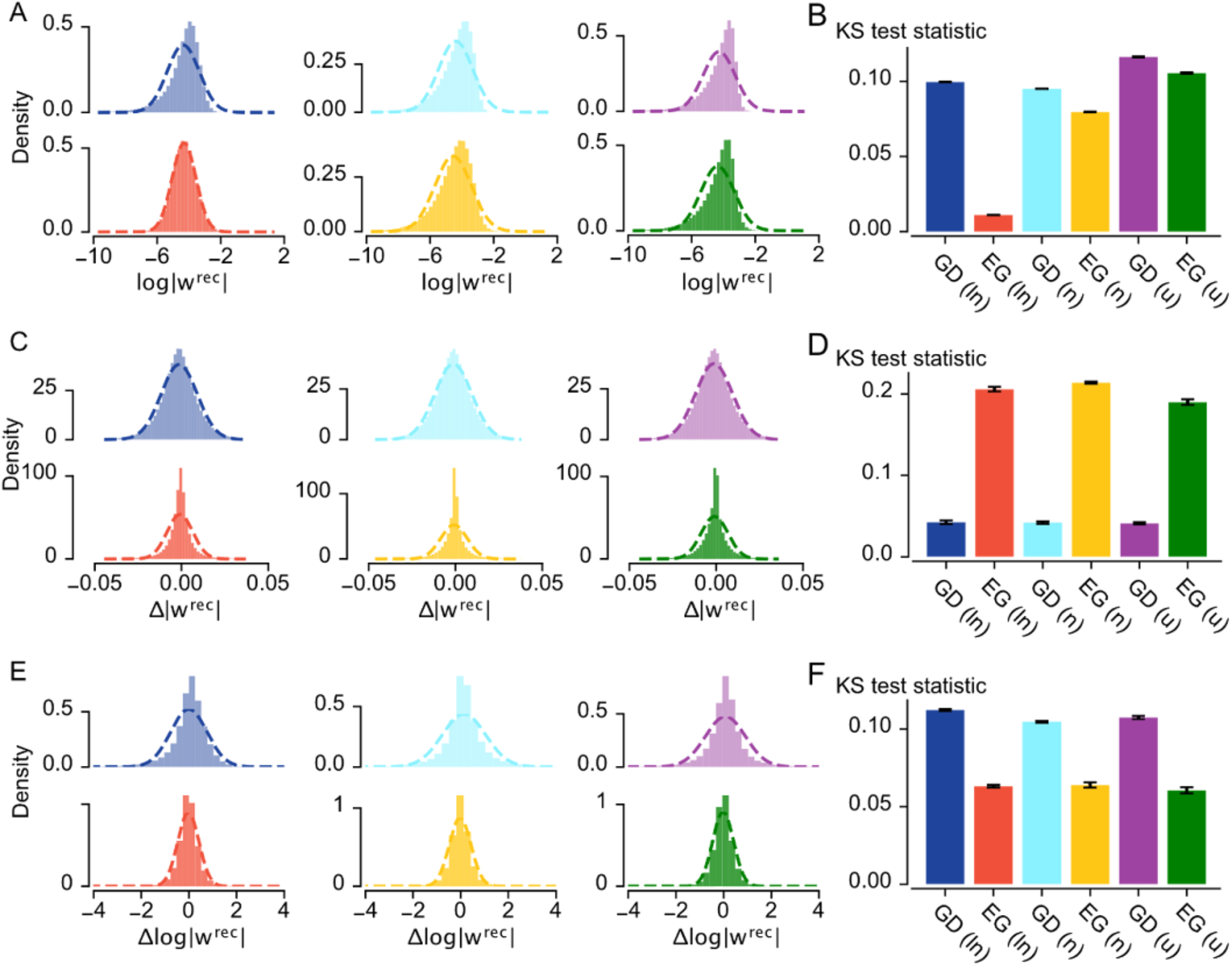
Dual space changes. Gaussian fits to final weights and weight changes for EG and GD for different weight initialisations (zero mean with equal variance). Log-normal: (ln); normal (n); uniform: (u): **a**| Final weights in the log space for GD (top row) and EG (bottom row) with (left to right) log-normal, normal, and uniform initialisations. **b**| Kolmogorov-Smirnov (KS) test statistic for standard normal fits of z-scored log final weights. EG with a log-normal distribution is much closer to a normal distribution in the log space, and EG for all initialisations is closer than GD. **c**| Same as **a**, but for weight changes. **d**| Same as **b**, but for weight changers. GD weight changes are consistently more normal across weight initialisations. **e**| Same as **a**, but for changes of log weights. **f**| Same as **b**, but for changes of log weights. EG weight changes in the log space are consistently more normal than those for GD (cf. **d**).

**Supplementary Figure 4:**
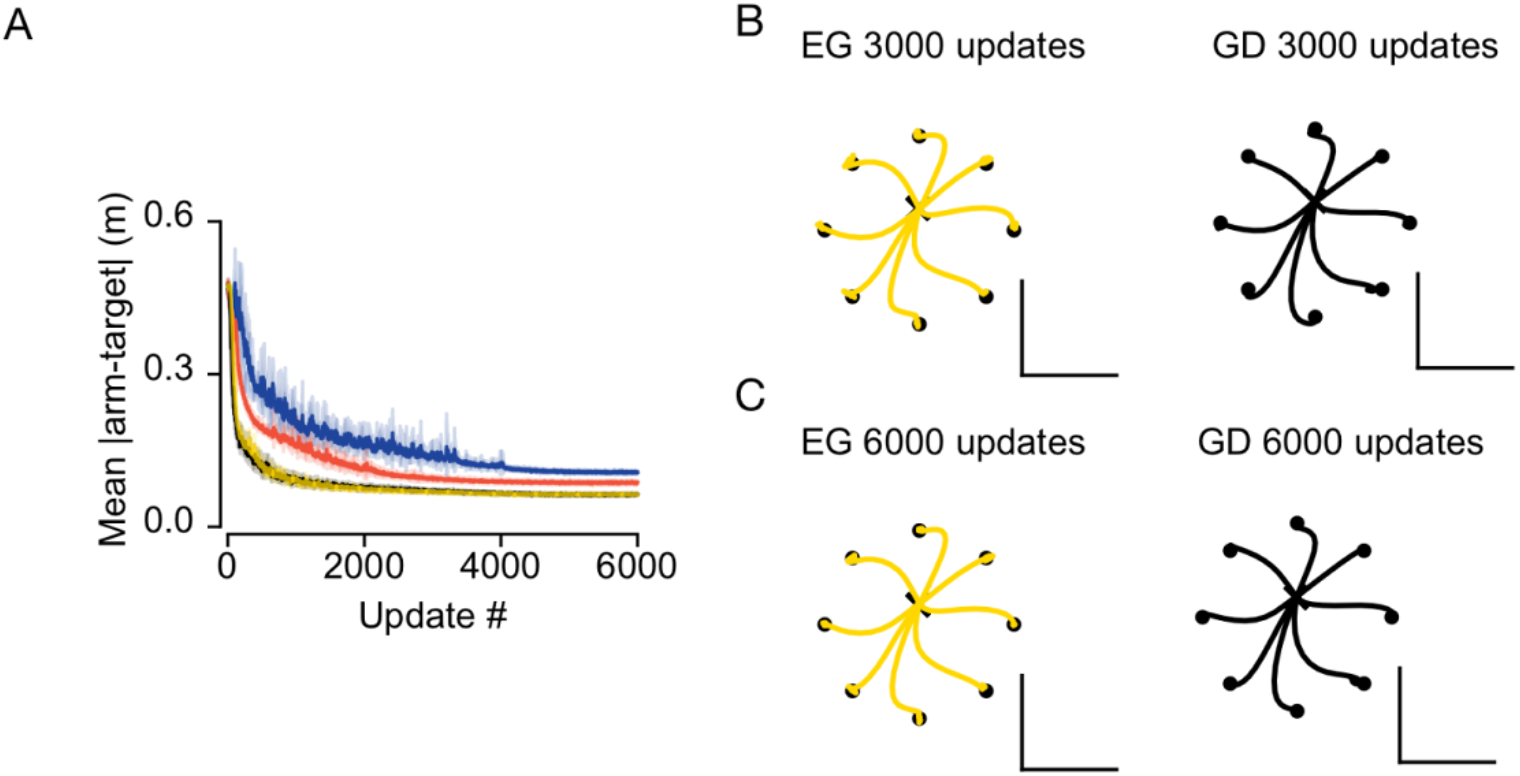
Continuous control. **a**| Learning curves for RNNs trained with EG (red, gold)or GD (blue, black). (From n=9 models, 500 distractors blue and red, 0 distractors gold and black).

## Notes

### Competing Interest Statement

The authors have declared no competing interest.

## References

Agrawal, K.K., Mondal, A.K., Ghosh, A., Richards, B., 2022. alpha-ReQ : Assessing Representation Quality in Self-Supervised Learning by measuring eigenspectrum decay, in: Koyejo, S., Mohamed, S., Agarwal, A., Belgrave, D., Cho, K., Oh, A. (Eds.), Advances in Neural Information Processing Systems. Curran Associates, Inc., pp. 17626–17638.

Allen Cell Types Database, 2015.

Bartunov, S., Santoro, A., Richards, B., Marris, L., Hinton, G.E., Lillicrap, T., 2018. Assessing the Scalability of Biologically-Motivated Deep Learning Algorithms and Architectures, in: Bengio, S., Wallach, H., Larochelle, H., Grauman, K., Cesa-Bianchi, N., Garnett, R. (Eds.), Advances in Neural Information Processing Systems. Curran Associates, Inc.

Bernstein, J., Zhao, J., Meister, M., Liu, M.-Y., Anandkumar, A., Yue, Y., 2020. Learning compositional functions via multiplicative weight updates. Adv. Neural Inf. Process. Syst. 33, 13319–13330.

Bicknell, B.A., Häusser, M., 2021. A synaptic learning rule for exploiting nonlinear dendritic computation. Neuron 109, 4001-4017.e10. 10.1016/j.neuron.2021.09.044

Bredenberg, C., Savin, C., 2023. Desiderata for normative models of synaptic plasticity. 10.48550/arXiv.2308.04988

Bregman, L.M., 1967. The relaxation method of finding the common point of convex sets and its application to the solution of problems in convex programming. USSR Comput. Math. Math. Phys. 7, 200–217.

Brown, T.B., Mann, B., Ryder, N., Subbiah, M., Kaplan, J., Dhariwal, P., Neelakantan, A., Shyam, P., Sastry, G., Askell, A., Agarwal, S., Herbert-Voss, A., Krueger, G., Henighan, T., Child, R., Ramesh, A., Ziegler, D.M., Wu, J., Winter, C., Hesse, C., Chen, M., Sigler, E., Litwin, M., Gray, S., Chess, B., Clark, J., Berner, C., McCandlish, S., Radford, A., Sutskever, I., Amodei, D., 2020. Language Models are Few-Shot Learners. 10.48550/ARXIV.2005.14165

Bubeck, S., 2015. Convex optimization: algorithms and complexity, Foundations and trends in machine learning. Now, Boston Delft.

Buzsáki, G., Mizuseki, K., 2014. The log-dynamic brain: how skewed distributions affect network operations. Nat. Rev. Neurosci. 15, 264–278. 10.1038/nrn3687

Codol, O., Michaels, J.A., Kashefi, M., Pruszynski, J.A., Gribble, P.L., 2024. MotorNet, a Python toolbox for controlling differentiable biomechanical effectors with artificial neural networks. Elife 12, RP88591.

Cornford, J., Kalajdzievski, D., Leite, M., Lamarquette, A., Kullmann, D., Richards, B., 2021. Learning to live with Dale’s principle: ANNs with separate excitatory and inhibitory units, in: ICLR 2021-9th International Conference on Learning Representations. ICLR.

Doerig, A., Sommers, R.P., Seeliger, K., Richards, B., Ismael, J., Lindsay, G.W., Kording, K.P., Konkle, T., Van Gerven, M.A., Kriegeskorte, N., others, 2023. The neuroconnectionist research programme. Nat. Rev. Neurosci. 24, 431–450.

Dorkenwald, S., Turner, N.L., Macrina, T., Lee, K., Lu, R., Wu, J., Bodor, A.L., Bleckert, A.A., Brittain, D., Kemnitz, N., Silversmith, W.M., Ih, D., Zung, J., Zlateski, A., Tartavull, I., Yu, S.-C., Popovych, S., Wong, W., Castro, M., Jordan, C.S., Wilson, A.M., Froudarakis, E., Buchanan, J., Takeno, M.M., Torres, R., Mahalingam, G., Collman, F., Schneider-Mizell, C.M., Bumbarger, D.J., Li, Y., Becker, L., Suckow, S., Reimer, J., Tolias, A.S., Macarico da Costa, N., Reid, R.C., Seung, H.S., 2022. Binary and analog variation of synapses between cortical pyramidal neurons. eLife 11, e76120. 10.7554/eLife.76120

Driscoll, L.N., Shenoy, K., Sussillo, D., 2024. Flexible multitask computation in recurrent networks utilizes shared dynamical motifs. Nat. Neurosci. 27, 1349–1363. 10.1038/s41593-024-01668-6

Eccles, J., 1976. From electrical to chemical transmission in the central nervous system: The closing address of the Sir Henry Dale Centennial Symposium Cambridge, 19 September 1975. Notes Rec. R. Soc. Lond. 30, 219–230. 10.1098/rsnr.1976.0015

Faust, T.E., Gunner, G., Schafer, D.P., 2021. Mechanisms governing activity-dependent synaptic pruning in the developing mammalian CNS. Nat. Rev. Neurosci. 22, 657–673. 10.1038/s41583-021-00507-y

Frankle, J., Carbin, M., 2018. The Lottery Ticket Hypothesis: Finding Sparse, Trainable Neural Networks. 10.48550/ARXIV.1803.03635

Gouwens, N.W., Berg, J., Feng, D., Sorensen, S.A., Zeng, H., Hawrylycz, M.J., Koch, C., Arkhipov, A., 2018. Systematic generation of biophysically detailed models for diverse cortical neuron types. Nat Commun 9, 710.

Gunasekar, S., Lee, J., Soudry, D., Srebro, N., 2018. Characterizing implicit bias in terms of optimization geometry, in: International Conference on Machine Learning. PMLR, pp. 1832–1841.

Gütig, R., Sompolinsky, H., 2006. The tempotron: a neuron that learns spike timing–based decisions. Nat Neurosci 9, 420–428.

Haber, A., Schneidman, E., 2022. The computational and learning benefits of Daleian neural networks. 10.48550/ARXIV.2210.05961

Hines, M., Carnevale, N., 1984. The NEURON Book. Cambridge University Press, Cambridge.

Hollander, M. A., Wolfe, D., Chicken, E., 2015. Nonparametric Statistical Methods, 1st ed, Wiley Series in Probability and Statistics. Wiley. 10.1002/9781119196037

Ikegaya, Y., Sasaki, T., Ishikawa, D., Honma, N., Tao, K., Takahashi, N., Minamisawa, G., Ujita, S., Matsuki, N., 2013. Interpyramid Spike Transmission Stabilizes the Sparseness of Recurrent Network Activity. Cereb. Cortex 23, 293–304. 10.1093/cercor/bhs006

Jahr, C.E., Stevens, C.F., 1990. Voltage dependence of NMDA-activated macroscopic conductances predicted by single-channel kinetics. J Neurosci 10, 3178–3182.

Khona, M., Chandra, S., Ma, J.J., Fiete, I., 2023. Winning the lottery with neural connectivity constraints: faster learning across cognitive tasks with spatially constrained sparse RNNs. 10.48550/arXiv.2207.03523

Kivinen, J., Warmuth, M.K., 1997. Exponentiated Gradient versus Gradient Descent for Linear Predictors. Inf. Comput. 132, 1–63. 10.1006/inco.1996.2612

Levenstein, D., Alvarez, V.A., Amarasingham, A., Azab, H., Chen, Z.S., Gerkin, R.C., Hasenstaub, A., Iyer, R., Jolivet, R.B., Marzen, S., Monaco, J.D., Prinz, A.A., Quraishi, S., Santamaria, F., Shivkumar, S., Singh, M.F., Traub, R., Rotstein, H.G., Nadim, F., Redish, A.D., 2023. On the role of theory and modeling in neuroscience. J. Neurosci. 43, 1074–1088. 10.1523/JNEUROSCI.1179-22.2022

Lillicrap, T.P., Santoro, A., Marris, L., Akerman, C.J., Hinton, G., 2020. Backpropagation and the brain. Nat. Rev. Neurosci. 21, 335–346. 10.1038/s41583-020-0277-3

Littlestone, N., 1988. Learning Quickly When Irrelevant Attributes Abound: A New Linear-Threshold Algorithm. Mach. Learn. 2, 285–318. 10.1023/A:1022869011914

Loewenstein, Y., Kuras, A., Rumpel, S., 2011. Multiplicative Dynamics Underlie the Emergence of the Log-Normal Distribution of Spine Sizes in the Neocortex In Vivo. J. Neurosci. 31, 9481–9488. 10.1523/JNEUROSCI.6130-10.2011

Mante, V., Sussillo, D., Shenoy, K.V., Newsome, W.T., 2013. Context-dependent computation by recurrent dynamics in prefrontal cortex. Nature 503, 78–84. 10.1038/nature12742

Maret, S., Faraguna, U., Nelson, A.B., Cirelli, C., Tononi, G., 2011. Sleep and waking modulate spine turnover in the adolescent mouse cortex. Nat. Neurosci. 14, 1418–1420. 10.1038/nn.2934

Mason-Williams, G., Dahlqvist, F., 2024. What Makes a Good Prune? Maximal Unstructured Pruning for Maximal Cosine Similarity, in: The Twelfth International Conference on Learning Representations.

Melander, J.B., Nayebi, A., Jongbloets, B.C., Fortin, D.A., Qin, M., Ganguli, S., Mao, T., Zhong, H., 2021. Distinct in vivo dynamics of excitatory synapses onto cortical pyramidal neurons and parvalbumin-positive interneurons. Cell Rep. 37, 109972. 10.1016/j.celrep.2021.109972

Molano-Mazon, M., Barbosa, J., Pastor-Ciurana, J., Fradera, M., Zhang, R.-Y., Forest, J., Del Pozo Lerida, J., Ji-An, L., Cueva, C.J., De La Rocha, J., Narain, D., Yang, G.R., 2022. NeuroGym: An open resource for developing and sharing neuroscience tasks. 10.31234/osf.io/aqc9n

Nemirovskij, A.S., Yudin, D.B., 1983. Problem complexity and method efficiency in optimization.

Omrani, M., Murnaghan, C.D., Pruszynski, J.A., Scott, S.H., 2016. Distributed task-specific processing of somatosensory feedback for voluntary motor control. eLife 5, e13141. 10.7554/eLife.13141

Pascanu, R., Mikolov, T., Bengio, Y., 2012. On the difficulty of training Recurrent Neural Networks. 10.48550/ARXIV.1211.5063

Pogodin, R., Cornford, J., Ghosh, A., Gidel, G., Lajoie, G., Richards, B., 2024. Synaptic Weight Distributions Depend on the Geometry of Plasticity.

Richards, B.A., Kording, K.P., 2023. The study of plasticity has always been about gradients. J. Physiol. 601, 3141–3149. 10.1113/JP282747

Rößler, N., Jungenitz, T., Sigler, A., Bird, A., Mittag, M., Rhee, J.S., Deller, T., Cuntz, H., Brose, N., Schwarzacher, S.W., Jedlicka, P., 2023. Skewed distribution of spines is independent of presynaptic transmitter release and synaptic plasticity, and emerges early during adult neurogenesis. Open Biol. 13, 230063. 10.1098/rsob.230063

Schwarz, J., Jayakumar, S.M., Pascanu, R., Latham, P.E., Teh, Y.W., 2021. Powerpropagation: A sparsity inducing weight reparameterisation. 10.48550/arXiv.2110.00296

Shalev-Shwartz, S., others, 2012. Online learning and online convex optimization. Found. Trends® Mach. Learn. 4, 107–194.

Song, S., Sjöström, P.J., Reigl, M., Nelson, S., Chklovskii, D.B., 2005. Highly Nonrandom Features of Synaptic Connectivity in Local Cortical Circuits. PLoS Biol. 3, e68. 10.1371/journal.pbio.0030068

Summerfield, C., 2022. Natural General Intelligence: How understanding the brain can help us build AI. Oxford university press.

Sun, H., Ahn, K., Thrampoulidis, C., Azizan, N., 2022. Mirror descent maximizes generalized margin and can be implemented efficiently. Adv. Neural Inf. Process. Syst. 35, 31089–31101.

Teramae, J., Fukai, T., 2014. Computational implications of lognormally distributed synaptic weights. Proc. IEEE 102, 500–512.

Todorov, E., Erez, T., Tassa, Y., 2012. MuJoCo: A physics engine for model-based control, in: 2012 IEEE/RSJ International Conference on Intelligent Robots and Systems. Presented at the 2012 IEEE/RSJ International Conference on Intelligent Robots and Systems (IROS 2012), IEEE, Vilamoura-Algarve, Portugal, pp. 5026–5033. 10.1109/IROS.2012.6386109

Van Rossum, M.C.W., Bi, G.Q., Turrigiano, G.G., 2000. Stable Hebbian Learning from Spike Timing-Dependent Plasticity. J. Neurosci. 20, 8812–8821. 10.1523/JNEUROSCI.20-23-08812.2000

Widrow, B., Hoff, M.E., 1988. Adaptive switching circuits, in: Neurocomputing: Foundations of Research. MIT Press, Cambridge, MA, USA, pp. 123–134.

Yang, G.R., Joglekar, M.R., Song, H.F., Newsome, W.T., Wang, X.-J., 2019. Task representations in neural networks trained to perform many cognitive tasks. Nat. Neurosci. 22, 297–306. 10.1038/s41593-018-0310-2

